# Prenatal environmental conditions underlie alternative reproductive tactics that drive the formation of a mixed-kin cooperative society

**DOI:** 10.1101/2021.02.10.430612

**Authors:** Shailee S. Shah, Dustin R. Rubenstein

## Abstract

Although animal societies often evolve due to limited natal dispersal that results in kin clustering and facilitates cooperation among relatives, many species form cooperative groups with low and variable kin structure. Groups in such mixed-kin societies often comprise residents and immigrants of the same sex that compete for breeding opportunities. To understand how such mixed-kin societies form despite their potential for social conflict, we investigated the environmental causes and subsequent fitness consequences of dispersal decisions in male cooperatively breeding superb starlings (*Lamprotornis superbus*) that live in a climatically unpredictable environment. We show that the two alternative reproductive tactics—natal dispersal or philopatry—exhibit reproductive tradeoffs resulting in equal lifetime inclusive fitness. The tactic an individual adopts is governed by the environment its parents experience prior to laying rather than the environment it experiences during its juvenile stage. When individuals adopt the tactic not predicted by early life environmental conditions, their fitness is reduced. Ultimately, climate-driven oscillating selection may help stabilize mixed-kin animal societies, despite their reduced kin structure and potential for social conflict.

## Main

Dispersal decisions that enable individuals to escape unfavorable environmental conditions or reduce inbreeding risk also influence ecological and evolutionary processes like range expansion, gene flow, and the formation of animal societies ^1,2^. Limited natal dispersal—the permanent movement of an individual from its natal to breeding site—plays a crucial role in social evolution by creating spatial clustering of kin that facilitates cooperative care of young by relatives ^3–5^. Although most animal societies are therefore characterized by groups with high kin structure, a surprising number of species form social groups with low kin structure ^6^ that consist of same-sex residents and immigrants, which often compete for breeding opportunities ^7–10^. Since indirect benefits alone cannot explain the evolution of these mixed-kin societies ^11–13^, direct reproductive or survival benefits may underlie their formation ^7,9,14^. Understanding how mixed-kin cooperative societies form and remain stable despite their reduced kin structure and potential for high social conflict will require examining the causes and lifetime fitness consequences of individual dispersal decisions to determine why some individuals remain in their natal groups with kin while others do not.

Dispersal decisions are influenced not only by aspects of the social or ecological environment that individuals experience during their lifetime after reaching nutritional independence ^15–17^, but also by how these factors affect their parents prior to birth and impact their development early-in-life (termed prenatal factors) ^18,19^. The influence of prenatal factors typically manifests via parental effects, modifications provided by the mother or father to the offspring during development that alter offspring phenotype ^20^. Parental effects may be particularly important in cooperatively breeding species ^21^, where parents can limit offspring dispersal by influencing helping behavior or promote dispersal to reduce kin competition ^19,22,23^. Although dispersal in most social species tends to be sex-biased ^24^, in many cooperative breeders, individuals of one sex can adopt either of two alternative reproductive tactics: natal dispersal or philopatry ^8,15^. Such sex-specific behavioral polyphenisms can exist either because (i) individuals face conditional constraints in adopting the tactic with higher fitness and must instead make “ the-best-of-a-bad-job” by choosing the tactic with lower fitness ^25,26^, or (ii) the relative fitness payoffs are environmentally-dependent such that both tactics can persist within a population when conditions fluctuate ^27,28^. Consistent with the “ best-of-a-bad-job” hypothesis, the fitness benefits of alternative reproductive tactics in cooperative breeders tend to be unequal in the short-term ^29–32^. Yet, few studies have compared the lifetime inclusive fitness outcomes of alternative reproductive tactics, especially in mixed-kin societies where they are most common ^8^. Thus, the evolutionary mechanism underlying the formation of mixed-kin societies remains largely unknown and can only be revealed by considering the lifetime inclusive fitness consequences of alternative reproductive tactics.

Here, we leveraged a 16-year longitudinal dataset from Kenya to examine how reproductive tactics in male cooperatively breeding superb starlings (*Lamprotornis superbus*) are influenced by ecological and social factors in both the prenatal and juvenile periods, and ultimately how these alternative tactics impact lifetime inclusive fitness. Superb starlings are plural cooperative breeders in which both sexes help ^33,34^, and although males tend to be more philopatric than females, these mixed-kin social groups have low but variable kin structure overall as well as within both sexes ^33,35^. While resident females never breed in their natal groups, both resident and immigrant males can acquire breeding status ^33^. These savanna-dwelling birds inhabit one of the most variable and unpredictable environments on earth ^36^, where annual fluctuation in seasonal rainfall governs insect prey availability ^37^. Rainfall is most variable during the dry season ^38^, and this variation has been shown to influence many aspects of superb starling social behavior and life-history. In particular, rainfall experienced by the parents (prenatal rainfall) influences offspring sex ratio ^34^, epigenetic modification of genes related to the avian stress response ^39^, and male reproductive decisions ^39^. Dry season rainfall experienced by individuals of both sexes also influences access to breeding opportunities ^40^ and helping behavior ^41^. In addition, the social environment (group size) also impacts superb starling fitness, both survival ^42^ and reproductive success ^38^.

To examine the environmental causes of alternative reproductive tactics in superb starlings, we first investigated how ecological and social factors during both the prenatal and juvenile stage influence male dispersal decisions. We predicted that under harsh prenatal environmental conditions (low rainfall), particularly for small groups with female-skewed sex ratios, parental effects may indirectly limit dispersal of male offspring by promoting helping behavior. Additionally, males that remain in their natal group would be more likely to provide alloparental care in their first year of life than males that ultimately disperse. Similarly, male superb starlings that experience harsh environmental conditions as juveniles, the life history stage when dispersal decisions are likely made ^33^, would be more likely to remain in their natal group in order avoid the costs of dispersal or reap the benefits of philopatry. Furthermore, a social environment conducive to higher survival (i.e., when in larger groups) and with greater access to reproductive opportunities (groups with female-skewed sex ratios) may also promote philopatry. Next, to understand the fitness consequences of these alternative reproductive tactics, we compared access to reproductive opportunities, reproductive success, lifetime inclusive fitness, and survivorship of resident and immigrant males. According to the best-of-a-bad-job hypothesis, we predicted that the two alternative tactics would result in unequal lifetime inclusive fitness. Alternatively, if both dispersal and philopatry maximize individual fitness, then the two tactics would have equal lifetime inclusive fitness but exhibit reproductive tradeoffs such that individuals would have lower lifetime fitness when they adopt the reproductive tactic not predicted by the prevailing environmental conditions. Ultimately, by linking the environmental causes and fitness consequences of alternative reproductive tactics in a cooperative breeder we will be able to determine how two fundamentally different reproductive phenotypes can persist and potentially give rise to mixed-kin, plural breeding societies.

## Results

### Patterns of male dispersal

Consistent with previous reports in this species ^33^, we found that males were the primary philopatric sex; only 59 of 198 males (30%) in our marked social groups for whom we had data on dispersal status (i.e., excluding individuals already present in the groups at the beginning of 2001 (dispersal status unknown) and individuals not seen past one year of age (presumed dispersed)) were immigrants (compared to 121 of 248 (49%) of females). However, among all males for whom we had complete life history data (N = 162), more than half the breeders (30 of 59; 51%) were immigrants.

### Ecological and social predictors of male dispersal

Next, we examined the role of ecological and social factors experienced by males during their juvenile stage (i.e., the period when dispersal occurs) as well as by their parents (i.e., prenatal) in influencing male dispersal decisions. We found that harsh prenatal ecological conditions were associated with limited dispersal such that males were less likely to disperse when they were born in years following low prenatal rainfall (Z = 2.03, P = 0.04, CI = 0.02-0.74; Fig. 1, Table 1). Since prenatal ecological conditions did not affect hatchling mass prior to fledging (see *SI*), this pattern was likely driven by dispersal, and not mortality. In contrast, the social environment as well as the ecological conditions experienced during the juvenile period had no significant impact on the likelihood of males dispersing (Table 1). Together, these results suggest that male dispersal is influenced more by (i) conditions experienced by their parents than the conditions they experience as juveniles and (ii) ecological rather than social factors. Finally, we also found that males that subsequently dispersed were less likely to act as alloparents in their first year of life than males that subsequently remained in their natal group (χ^2^_1_ = 7.63, P = 0.006).

**Table 1.** Factors affecting dispersal decisions of natal male superb starlings. Results of a generalized linear mixed model with a binomial response (dispersed vs. remained) for natal males (N = 185). Since sex ratio and group size are highly correlated within a year, we only included the former in the final model. Breeding season was included as a random effect. Random effects of social group, father ID, mother ID, and season (short vs. long rains) had variance components equal to zero, and were thus excluded from the final model to facilitate the computation of 95% confidence intervals.

| Effect | Estimate | Std. Error | Z value | 95% CI |  | p-value |
| --- | --- | --- | --- | --- | --- | --- |
| Intercept | -0.97 | 0.18 | -5.45 | -1.36 | -0.60 | <0.001 |
| Prenatal rainfall | 0.36 | 0.18 | 2.03 | 0.02 | 0.74 | 0.04 |
| Prenatal sex ratio | -0.13 | 0.20 | -0.68 | -0.54 | 0.24 | 0.50 |
| Prenatal group size | 0.05 | 0.19 | 0.27 | -0.34 | 0.44 | 0.78 |
| Juvenile rainfall | -0.12 | 0.17 | -0.69 | -0.48 | 0.24 | 0.49 |

**Figure 1.**
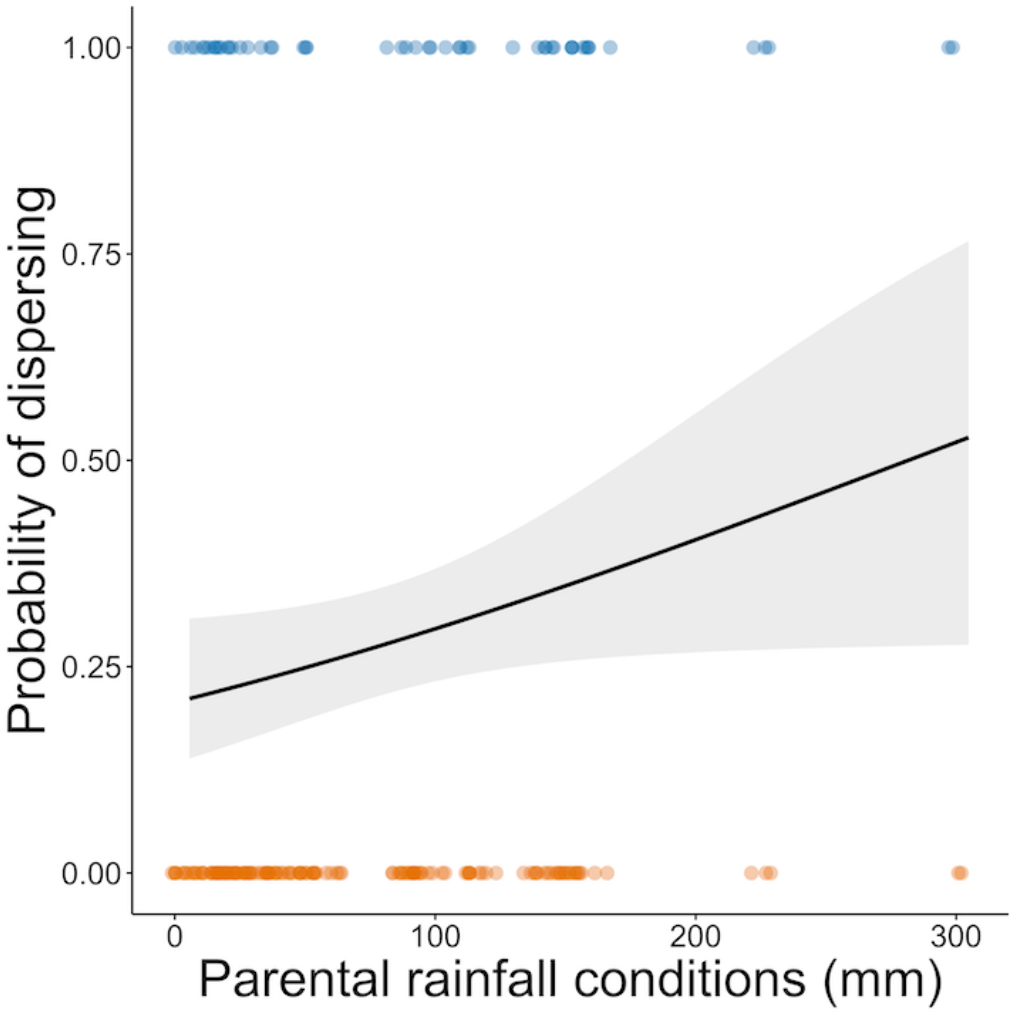
Effect of prenatal rainfall on the probability of male superb starlings dispersing from their natal groups. Males were more likely to disperse when born following periods of higher prenatal rainfall (N = 185, P = 0.04). Model estimate (solid line) is bounded by the 95% confidence interval (shaded areas). Circles indicate raw data (blue = dispersed, orange = remained).

### Fitness consequences of male dispersal decisions

We then explored the fitness consequences of the different male dispersal decisions. More than half the males in the population (85 of 152 males; 56%) failed to accrue any inclusive fitness in their lifetimes (Fig. S3) ^33^. Residents unsurprisingly accrued higher indirect fitness than immigrants (Fig. 2C, U = 2103, P = 0.02), who were unlikely to accrue meaningful indirect fitness in their natal group prior to dispersal (see *SI*). In contrast, resident and immigrant males had similar direct and inclusive lifetime fitness (Fig. 2A-B; inclusive: U = 2337, P = 0.22; direct: U = 2701, P = 0.59). While this analysis does not quantify indirect fitness accrued by immigrants in their natal group prior to dispersing, natal males that dispersed accrued significantly lower fitness in their first year than residents, suggesting that immigrants into our study population also did not likely accrue considerable fitness prior to dispersing (*see SI*). Nonetheless, to account for the possibility that immigrant males accrued at least some indirect fitness in their natal group prior to dispersal, we repeated our analysis after excluding fitness accrued by residents in their first year of life. We found qualitatively similar results, with residents and immigrants still having similar direct and inclusive fitness, as well as similar indirect fitness (see *SI*).

**Figure 2.**
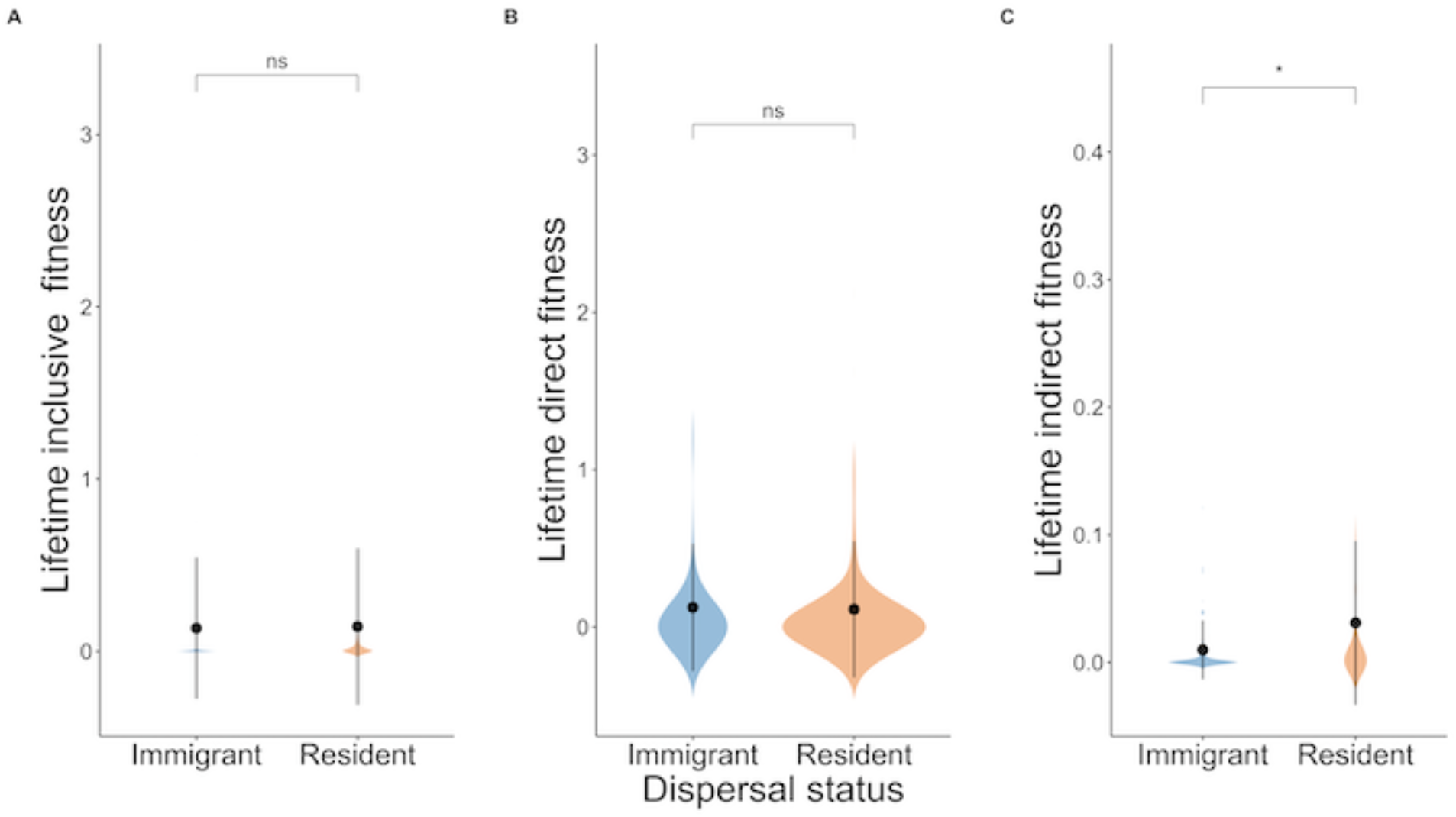
Mean lifetime fitness of immigrant and resident male superb starlings. Immigrant and resident males (N_Imm_ = 59, N_Res_ = 99) have equal lifetime (A) inclusive (P = 0.22) and (B) direct fitness (P = 0.59), though residents have higher (C) indirect fitness (P = 0.02). Black dots represent means and error bars denote standard deviation. Shaded areas are kernel probability densities illustrating the distribution of the data. Asterisks indicate significance (ns: P > 0.05; *: P < 0.05).

Although the survival likelihood of immigrants after successful dispersal is not significantly different from that of residents (Z = 0.06, P = 0.95, CI = -0.36-0.38) (Fig. S4), our analysis did not account for any potential mortality of immigrants during dispersal. While we expect the mortality of immigrants to be low because there are no floaters in this system^33^, to account for potential immigrant mortality during dispersal, we ran a sensitivity analysis that added immigrants with zero fitness in 10% increments to the analyses of direct, indirect, and inclusive lifetime fitness. Our sensitivity analysis found that the mortality threshold for immigrants at which direct fitness would be higher in residents than immigrants was 70% (Table S1). Similarly, the mortality threshold for immigrants at which inclusive fitness would be higher in residents than immigrants ranged from 20% to 50%, depending on whether first-year resident males were included in the analysis (Table S1). Together, these results indicate that resident and immigrant males have similar lifetime inclusive fitness, even when accounting for the potential of mortality during dispersal.

Despite similar lifetime inclusive fitness, immigrants were more likely than residents to breed at least once in their lifetimes (χ^2^_1_ = 7.39, P = 0.006), had higher lifetime breeding effort (the number of breeding attempts as a proportion of an individual’s adult lifespan) (U = 3869, P = 0.006), and began breeding at a younger age (U = 265.5, P = 0.01). In contrast, residents that did breed had higher nest success than immigrant breeders (Z = 2.46, P = 0.01, CI = 0.27-2.34). In addition, both lifetime breeding effort and dispersal status affected a male’s likelihood of accruing some nonzero inclusive fitness during his lifetime (Fig. 3A, Table 2). Among males with some lifetime inclusive fitness, residents accrued higher inclusive fitness than immigrants with the same amount of lifetime breeding effort (Fig. 3B, Table 2).

**Table 2.** Effects of lifetime breeding effort and dispersal status on lifetime inclusive fitness of male superb starlings. Results of generalized linear mixed models with a (A) binomial (zero vs. nonzero) (N = 152) and (B) continuous, positive (N = 67) response variable for lifetime inclusive fitness of males. Social group was included as a random effect in the first model, but since it explained none of the variance and hindered the calculation of 95% confidence intervals for the fixed effects, we removed it from the second model.

| Effect | Estimate | Std. Error | Z value | 95% CI |  | p-value |
| --- | --- | --- | --- | --- | --- | --- |
| Intercept | -1.10 | 0.35 | -3.14 | -1.86 | -0.44 | 0.002 |
| Lifetime breeding effort | 1.03 | 0.31 | 3.28 | 0.46 | 1.72 | 0.001 |
| Dispersal status | 0.78 | 0.39 | 2.02 | 0.01 | 1.56 | 0.04 |

(B) Response variable: continuous, positive lifetime inclusive fitness
| Effect | Estimate | Std. Error | t value | 95% CI |  | p-value |
| --- | --- | --- | --- | --- | --- | --- |
| Intercept | 0.02 | 0.01 | 2.04 | 0.01 | 0.06 | 0.04 |
| Lifetime breeding effort | 0.55 | 0.15 | 3.73 | 0.33 | 0.95 | <0.001 |
| Dispersal status | 0.04 | 0.02 | 2.27 | -0.00 | 0.08 | 0.03 |

**Figure 3.**
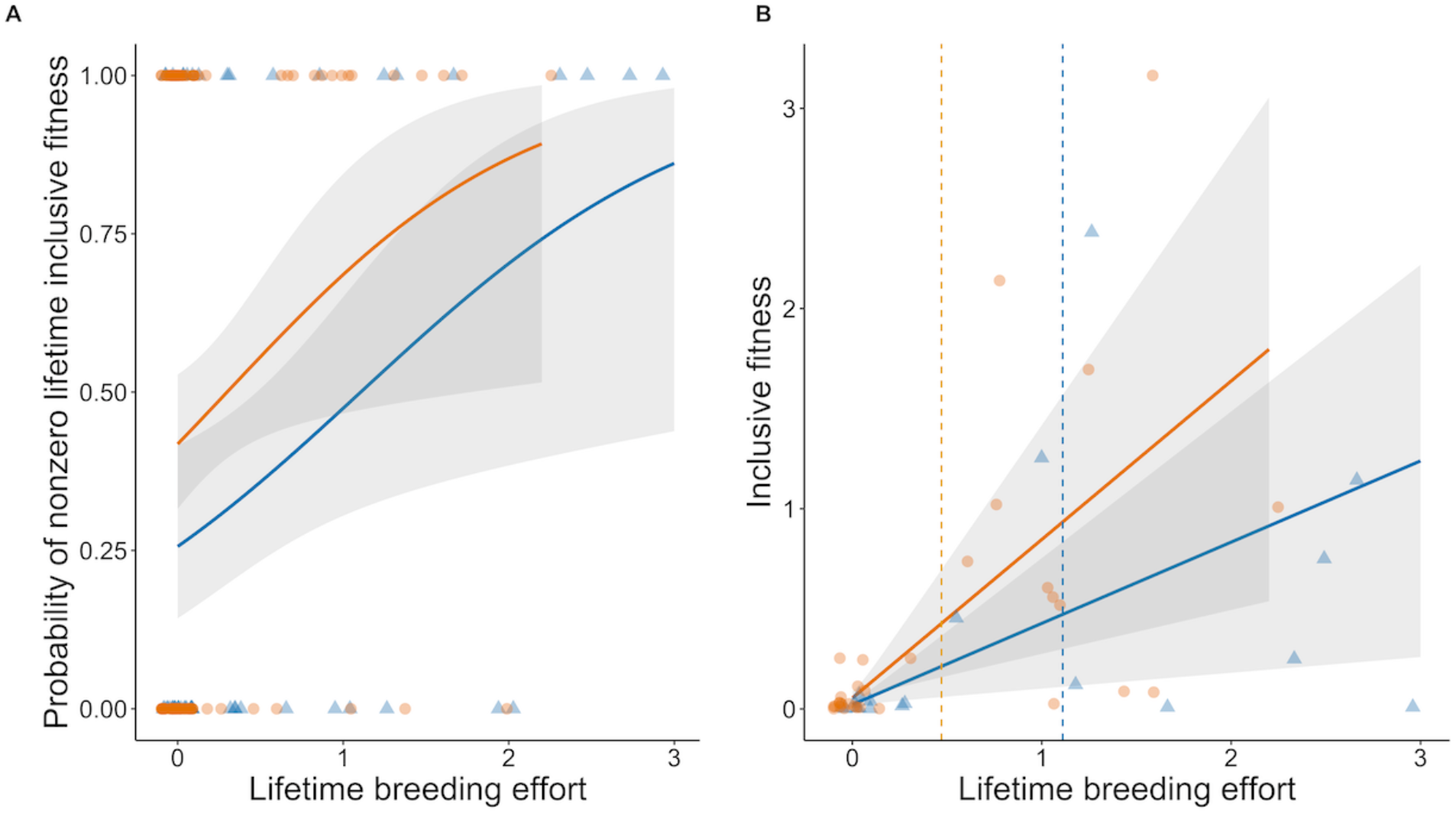
Effect of reproductive tradeoffs on lifetime inclusive fitness of male superb starlings. The effect of lifetime breeding effort and dispersal status (immigrants = blue, triangles; residents = orange, circles) on (A) the probability of accruing any nonzero lifetime inclusive fitness (N = 152), and (B) the value of nonzero lifetime inclusive fitness (N = 67). For visualization purposes, we excluded one individual with lifetime breeding effort greater than three standard deviations above the mean; this individual was included in the statistical analysis, though excluding it did not alter the results. Lifetime breeding effort (i.e., the number of breeding attempts as a proportion of an individual’s adult lifespan) (P = 0.001) and dispersal status (P = 0.04) affected a male’s likelihood of accruing some nonzero inclusive fitness during his lifetime. Among males with some lifetime inclusive fitness, residents accrued higher inclusive fitness than immigrants for the same amount of lifetime breeding effort (P = 0.04). Model estimates (solid lines) are bound by 95% confidence intervals (shaded areas). Points indicate raw data. Dashed vertical lines indicate mean lifetime breeding effort.

Finally, we assessed whether there were reproductive tradeoffs for individuals that adopted the tactic not predicted by prenatal ecological conditions. Since we did not have precise rainfall data from the immigrants’ natal sites (though rainfall is highly spatially correlated and temporally synchronous within the dispersal radius of our study site; see *SI*), we used a categorical measure of prenatal rainfall based on the mean long-term pre-breeding rainfall at our study site (see *Methods* and *SI*). Consistent with the idea of reproductive tradeoffs and the hypothesis that the fitness consequences of alternative reproductive tactics depend on prenatal environmental conditions, we found that immigrant males were more likely to accrue nonzero lifetime inclusive fitness when born following periods of high prenatal rainfall, but that residents were more likely to accrue nonzero lifetime inclusive fitness when born following periods of low prenatal rainfall (Fig. 4, Table 3).

**Table 3.** Effects of male superb starlings adopting the tactic not predicted by the prevailing prenatal ecological conditions on the likelihood of gaining nonzero inclusive fitness during their lifetimes. Results of a generalized linear mixed model with a binomial response (zero vs. nonzero lifetime inclusive fitness) variable for males (N = 152). Social group was included as a random effect.

| Effect | Estimate | Std. Error | Z value | 95% CI |  | p-value |
| --- | --- | --- | --- | --- | --- | --- |
| Intercept | -0.10 | 0.46 | -0.23 | -1.03 | 0.80 | 0.82 |
| Categorical prenatal rainfall | -0.63 | 0.59 | -1.08 | -0.80 | 0.52 | 0.28 |
| Dispersal status | -0.84 | 0.64 | -1.31 | -2.13 | 0.40 | 0.19 |
| Categorical prenatal rainfall x Dispersal status | 1.74 | 0.77 | 2.25 | 0.24 | 3.29 | 0.02 |

**Figure 4.**
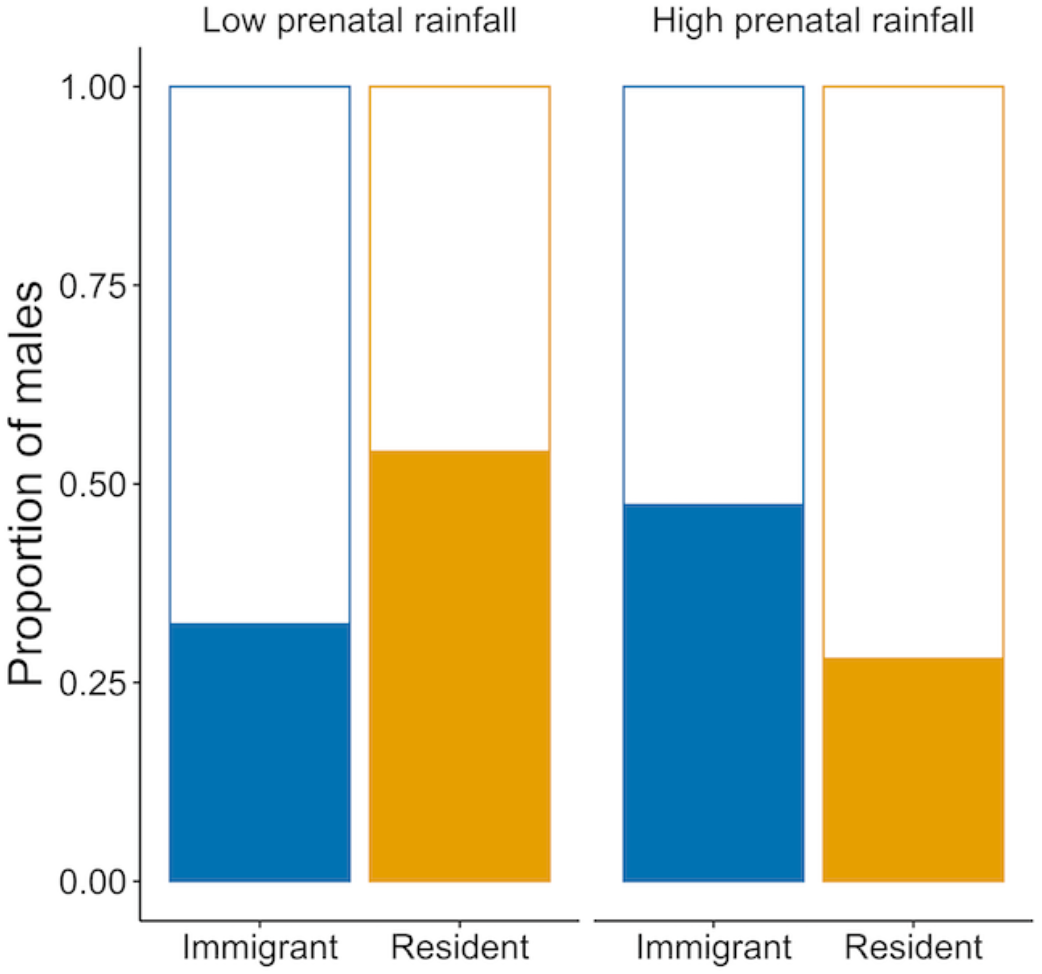
Fitness consequences of male superb starlings adopting the tactic not predicted by the prevailing prenatal environmental conditions. The effect of prenatal rainfall and dispersal status (immigrants = blue, residents = orange) on the probability of accruing any nonzero lifetime inclusive fitness (fill = nonzero fitness, no fill = zero fitness) (N = 152). Immigrants were more likely to accrue nonzero lifetime inclusive fitness when born following periods of high prenatal rainfall, whereas residents were more likely to accrue nonzero lifetime inclusive fitness when born following periods of low prenatal rainfall (P = 0.02).

## Discussion

Although mixed-kin cooperative groups characterized by low relatedness are surprisingly common (45% of all cooperatively breeding birds)^6^, it remains unclear how these societies arise and are maintained given the potential for high social conflict among unrelated group members competing for reproductive opportunities. We examined the environmental causes and inclusive fitness consequences of dispersal decisions in male superb starlings living in a climatically variable and unpredictable savanna environment ^36,42^, for whom philopatry and dispersal can both be pathways to reproductive opportunities ^33^. We found that dispersal in males is influenced by their parents’ prenatal environment rather than the environment that the males experience as juveniles, suggesting that prenatal parental effects play an important role in governing dispersal decisions. Ultimately, natal dispersal and philopatry had equal lifetime inclusive fitness outcomes, meaning that they are two equal, alternative male reproductive tactics. Yet, the two tactics showed reproductive tradeoffs—immigrants had greater access to reproductive opportunities via the acquisition of dominant breeding positions, whereas residents that bred had higher reproductive success. When males adopted a tactic not predicted by prevailing prenatal environmental conditions, their lifetime inclusive fitness was reduced. Together, these results suggest that environmental variability experienced early in life may help maintain a reproductive polyphenism with equal lifetime inclusive fitness, which ultimately enables the formation and persistence of a mixed-kin cooperative society in an unpredictable environment.

Unlike in other cooperatively breeding species where conditions experienced during the juvenile stage shape dispersal decisions ^15,16,43^, male dispersal in superb starlings was only governed by prenatal ecological conditions. Although we were unable to directly distinguish between dispersal and morality, several lines of evidence—including the modal age at which natal males disappear from and modal age of immigrants into our study population, the age at which males reach sexual maturity, the close match in the proportions of dispersers and immigrants, the increased likelihood of dispersal following benign ecological conditions, no effect of prenatal ecological conditions on hatchling mass prior to fledging, and the high likelihood of post-fledgling survival from year-to-year ^42^—strongly suggest that the pattern we observed is driven by dispersal, and not mortality (see *Methods and SI* for further details). The effect of prenatal conditions on dispersal suggests that parental effects influence the dispersal phenotype of male superb starlings, a result that has been suggested previously for male reproductive opportunities in this species ^39^. Parental effects, which can result from physiological or molecular modifications to offspring during development ^20^, may be particularly important in cooperatively breeding species ^21^ and species inhabiting unpredictable environments ^44^. While the specific physiological mechanisms of prenatal parental effects on dispersal remain unknown for this species, previous studies have demonstrated an effect of maternal condition on offspring sex allocation, maternal investment in eggs, and DNA methylation in superb starlings ^34,39^ and other vertebrates ^23,45,46^. Studies of two other avian species, western bluebirds (*Sialia mexicana*) and great tits (*Parus major*), found an effect of maternal androgen deposition on offspring dispersal in response to ecological conditions ^18,19^. Viviparous lizards (*Lacerta vivipara*) show a similar effect ^46^, and maternal hormones have also been shown to increase helping behavior in subordinate female meerkats (*Suricata suricatta*) ^22^. Alternatively, conditions experienced by parents may affect their behavior towards their offspring after birth, such that dispersal is driven purely by behavioral interactions prior to offspring becoming self-sufficient ^47,48^. Further work in this and other cooperatively breeding species should examine the mechanism by which environmentally-mediated parental effects might influence future reproductive tactics adopted by offspring.

Parental effects can be selfish, to manipulate offspring phenotype in order to increase parental fitness, or anticipatory, to maximize offspring fitness ^21,49^. We know that offspring sex ratio in superb starlings is male-biased when mothers are in poorer body condition following harsher prenatal conditions, a pattern that suggests that females maximize their inclusive fitness in an unpredictable environment by investing in the sex with lower fitness variance ^34^. However, our results suggest that mothers could also be influencing cooperative behavior in their offspring because males that participated in alloparental care in their first year of life were subsequently more likely to remain in their natal group. Male superb starlings are more likely to become alloparents and help more than females ^38^, and alloparents buffer the detrimental effects of harsh environmental conditions on reproductive success in this species ^41^. Thus, prenatal parental effects promoting alloparental behavior following harsher prenatal conditions might serve to selfishly increase the parents’ short-term future reproductive success and indirectly limit dispersal of male offspring ^22^. Although males born following benign prenatal conditions could, conversely, be more likely to disperse so as to reduce kin competition, this seems less probable because kin competition has actually been shown to decrease under benign environmental conditions in other cooperatively breeding species ^50,51^. Additionally, our results are consistent with the hypothesis that prenatal parental effects on dispersal in males are anticipatory because when males adopted the tactic predicted by the prevailing prenatal ecological conditions, they were more likely to accrue some inclusive fitness in their lifetimes. Since more than half of superb starling males fail to accrue any fitness in their lifetimes, largely due to high nest predation pressure ^33^, securing any amount of nonzero lifetime fitness is crucial in this species, an idea consistent with the bet-hedging hypothesis proposed for this ^38^ and other social species ^52^. Males born following harsh prenatal conditions thus appear to benefit from avoiding the costs of dispersal and remaining in their natal group ^17,53^, whereas males born following benign conditions might be better able to cope with the costs of dispersal and maximize their fitness by immigrating into another social group ^31^.

In contrast to other studies in kin-only cooperatively breeding societies ^29,30,54–57^, we found that the two reproductive tactics adopted by males—natal dispersal and philopatry—have equal lifetime inclusive fitness in a mixed-kin cooperative society. Although resident males had significantly higher lifetime indirect reproductive success than immigrants, an unsurprising result given the kin structuring among males in superb starling social groups ^33^, this was more than balanced out by the relatively larger contribution of lifetime direct fitness, even when accounting for potential mortality during dispersal. While it was not logistically possible to compare the fitness of resident and dispersing males born to the same social group, our approach of comparing resident to immigrant males has proven informative in other avian studies ^58^. While we do not have an estimate of dispersal mortality in this species, we have never observed floaters in our study area, suggesting that individuals move from their natal group to their non-natal group over a relatively short time period and likely do not incur significant mortality in transit ^33^. Following successful dispersal, our results show that the survival likelihood of immigrants is similar to that of resident males. Furthermore, natal males that subsequently remain in their natal group accrue significantly higher indirect fitness in their first year of life, suggesting that immigrants are unlikely to accrue considerable indirect fitness in their natal group prior to dispersing. Yet, when even accounting for this possibility, immigrant and resident males still had similar lifetime inclusive fitness. Similar variation in alloparental effort and indirect fitness in relation to dispersal tactics has been shown in other cooperatively breeding species ^58,59^.

Finally, we also show that the equal lifetime inclusive fitness outcomes of the two dispersal tactics are due to a reproductive tradeoff, an idea that has been proposed theoretically ^28,60^ but rarely tested empirically. Although immigrant superb starlings had greater access to reproductive opportunities via the acquisition of dominant breeding positions with the group, residents that acquired breeding positions were more likely to successfully fledge young. Residents can therefore afford to have lower access to breeding opportunities over their lifetimes since they have higher nest success. This higher nest success is likely due to greater alloparental care at nests of resident males who have more kin in the group to act as helpers than do immigrants ^61–63^. Thus, for superb starling males, philopatry leads to lower reproductive quantity but higher quality, whereas dispersal results in higher reproductive quantity but lower quality. Since immigrants and residents are equally likely to survive from one breeding season to the next, these differences in reproductive quality and quantity are unrelated to longevity ^29^. Together these results suggest that the two alternative dispersal tactics are equivalent in terms of lifetime inclusive fitness and refute the hypothesis that remaining in the natal group—the tactic favored following harsh prenatal conditions—is simply making “ the-best-of-a-bad-job” ^25,26^. The coexistence of resident and immigrant males in cooperative social groups has been shown to be similarly facilitated by equal reproductive rates in spotted hyenas (*Crocuta crocuta*) ^8^ and inclusive fitness at different age stages in dwarf mongooses (*Helogale parvula*) ^64^, both species that live in mixed-kin societies. However, to the best of our knowledge, ours is the first study in a cooperatively breeding vertebrate to demonstrate equal lifetime inclusive fitness outcomes of alternative dispersal tactics.

This behavioral polyphenism in reproductive tactics is likely maintained by a flexible response to high and unpredictable environmental variability ^27^. A temporally variable and unpredictable environment has been shown both theoretically ^28^ and empirically ^65^ to generate conditional strategies that result in a developmental switch between alternative tactics. If alternative reproductive tactics have a strong genetic (or perhaps epigenetic) basis ^66,67^, environmental variability can reverse the selective differential between the two tactics from one year to the next, resulting in alternative tactics with equal fitness outcomes ^28,68^. The savanna habitat inhabited by superb starlings has high temporal ecological variability ^36,38^ that may allow both dispersal tactics to persist, resulting in the formation of mixed-kin cooperative groups.

Interestingly, spotted hyenas ^69^ and dwarf mongooses ^64^ experience the same unpredictable African savanna environments as superb starlings, suggesting that environmental uncertainty—and perhaps oscillating selection pressures more generally—maintains alternative dispersal tactics and leads to the formation of and/or stabilizes societies with low kin structure. Oscillating selection may be particularly important in arid and semi-arid environments where climatic variability is high ^68^, and may help explain the evolution of both mixed-kin societies and plural breeding more generally. Indeed, many other plural cooperatively breeding birds across the globe live in harsh arid and semi-arid environments characterized by unpredictable variation in rainfall and food resources ^13,70–77^. While we do not yet know whether dispersal has a genetic basis in superb starlings—though there are indications of a potential epigenetic basis ^39^—our results suggest that the two tactics face oscillating selection since the relative lifetime inclusive fitness of the two tactics fluctuates in response to prenatal environmental conditions. Ultimately, identifying the mechanism by which prenatal parental effects lead to developmental differences that determine male reproductive tactics will be important for understanding the role that environmentally-driven selection pressures play in shaping behavioral polyphenisms in this and other social species.

In summary, we have shown that natal dispersal and philopatry in male cooperatively breeding superb starlings represent two alternative reproductive tactics with equal lifetime inclusive fitness. The tactics are likely mediated by parental effects during development and are maintained by oscillating selection pressures characteristic of their variable and unpredictable savanna environment. Our study suggests a direct link between environmental uncertainty, behavioral polyphenism, and the evolution of mixed-kin animal societies that cannot be explained by indirect fitness benefits alone. Our work also underscores the importance of prenatal environmental conditions and parental effects in determining offspring phenotype, especially in cooperative societies where early life conditions have direct implications on the future fitness of both parents and offspring. Moreover, the direct benefits derived from environmental selective pressures appear to play a significant role in the evolution and maintenance of cooperative societies, alongside or in the absence of kin selection. Ultimately, understanding how fluctuation in early life environmental conditions helps mediate reproductive tradeoffs and lifetime fitness is critical in an era of rapid anthropogenic climate change because climatic uncertainty is only likely to increase across much of the globe for the foreseeable future.

## Materials and Methods

### Data collection

Nine superb starling social groups have been monitored continuously since 2001 at the Mpala Research Centre, Kenya (0° 17’N 37° 52’E) ^33,78^. The study population is distributed over a 9 km extent from North to South. Groups (mean size ± SD = 22 ± 12) defend stable territories year-round and consist of breeding (mean ± SD = 2.70 ± 1.49 pairs per group) and non-breeding individuals, some of whom act as alloparents that guard and/or provision young ^33,41^. Birds breed twice a year during the long (March-June) and short rains (October-November) ^33^. We used data from the beginning of the 2001 short rains breeding season through the end of the 2017 long rains breeding season (N = 33 breeding seasons over 16 years), since not all birds in the study population were banded in the first breeding season. Limited focal observation data for nests in 2001 short rains was accounted for in our analyses (see *Data Analysis*).

Birds were banded with a unique combination of colored leg bands and a numbered metal ring. Hatchlings were banded in the nest; all other individuals were captured in baited pull-string traps or mist nets and banded after fledging from the nest ^78^. If banded post-fledgling, age was assessed via eye color as fledgling (black eyes), sub-adult (less than one year of age; cloudy eyes), or adult (one year of age or older; white eyes) ^42^. Dispersal status (i.e., resident or immigrant) was assigned based on age at banding and genetic parentage. Residents were either banded as hatchlings in the nest or as juveniles whose parents were genetically identified as members of the same group using 15 microsatellite markers ^79^. Immigrants were defined as those banded as juveniles or older whose parents were not genetically identified as belonging to the same group. We also used the same microsatellite markers to estimate (i) parentage and direct reproductive success in Cervus v3.0 ^80^ using methods described previously for this species ^40,81–83^ and (ii) pairwise relatedness ^84^ between all individuals with the R package *related* ^85^. Sex was determined genetically ^86^ as previously described for this species ^78^.

We performed daily nest searches throughout the breeding season. Active nests were observed with a spotting scope for 60-120 min per observation period (total observation time per nest: mean ± SD = 314.33 ± 248.82 min). All superb starlings within 30 m of the nest were identified, and their times of arrival and departure were recorded ^83^. Parents are the primary nest builders, and only the mother incubates the eggs ^83^. All other members of the social group seen visiting or guarding the nest were categorized as alloparents ^41^.

Census data were used to estimate group size and sex ratio (calculated by dividing the number of males by the sum of number of males and immigrant females in the group to estimate potential mate competition. Natal females were excluded since they never breed and are thus not viable mates for males in the group). Groups were opportunistically censused year-round and each individual marked as either present or absent in its social group twice a year in six-month increments. Individuals not seen for five or more breeding seasons (i.e. 2.5 years) were assumed to be dead ^35,42^. We inferred that the dispersal window for males is between fledging and around one year of age (age of sexual maturity) using four lines of evidence: (i) the modal age at which males disappear from their natal group; (ii) the categorical age of immigrant males dispersing into the study population; (iii) the likelihood of males being detected in the census in their first year of life; and (iv) the minimum age of first breeding by resident males (see *SI* for details). If a male was observed in its natal group after one year of age, we classified it as a resident; if not, we classified it as having dispersed. Although we could not distinguish between dispersal and mortality, the positive relationship between the likelihood of dispersal and benign ecological conditions (see *Results*) suggests that these males did indeed disperse since we would have expected a negative relationship if mortality were higher following harsh ecological conditions. In addition, we found no variation in hatchling mass before fledging (measured at 10 days of age) with variation in ecological conditions, further suggesting that this pattern is driven by dispersal not mortality (see *SI*). Following the same lines of evidence, immigrant males were assumed to have entered the group at about one year of age (see *SI*).

We considered the period between end of the breeding season when a male is born to the beginning of the breeding season one year later (i.e., its juvenile stage) as its dispersal window (see *SI* for more details). Ecological and social conditions experienced by juveniles in this period were termed juvenile stage conditions. In contrast, ecological and social conditions experienced by parents in the pre-breeding period preceding the breeding season of an individual’s birth were termed prenatal conditions. A breeding season was defined as the period between two weeks before the first nest and two weeks after the last nest of the season. The period between breeding seasons was termed the pre-breeding period. Daily rainfall was measured using an automated Hydrological Services TB3 Tipping Bucket Rain Gauge at Mpala Research Centre ^87^, supplemented by a manual gauge at the same location when the automated gauge failed ^38^. To calculate prenatal and juvenile period rainfall, we summed the rainfall within the prenatal and juvenile stages of an individual’s lifetime, as defined above.

### Data analysis

Our dataset comprises individuals that are (i) known to have been born in the groups (“ natal”), (ii) known to have immigrated into the group (“ immigrant”), and (iii) were already part of the group in 2001 when the study population was banded. All individuals are either known to have disappeared (classified as either “ dead” or “ dispersed” depending on age at disappearance) or were still present in the study population in 2017. Natal individuals that did not disperse were termed “ residents”. Thus, individuals with “ known complete life histories” are natal and immigrant individuals who were classified as dead.

Using data from individuals with known complete life histories as well as natal individuals that are still alive and present in the study population past one year of age (N_M_ =198, N_F_ = 248), we first examined patterns of male dispersal by quantifying the proportion of immigrant adult males and females in the study population. We then determined how breeding opportunities were shared between immigrants and residents by calculating the proportion of immigrant breeders of both sexes in the study population using data from breeders with known complete life histories (N_M_ = 59, N_F_ = 121).

Next, using data from all natal males—including those classified as dispersed or dead as well as those still present in the study population past the age of 1 year in 2017—we investigated the effect of ecological and social conditions during the prenatal and juvenile stages on the likelihood of a male dispersing. We built a generalized linear mixed model (GLMM) with a binomial error structure and “ logit” link function. We used the binomial response of reproductive tactic (1 = dispersed, 0 = remained) as the dependent variable (N_dispersed_ = 52, N_remained_ = 133). Fixed effects, which were standardized using z-scores ^88^, included prenatal and juvenile stage rainfall, prenatal group size, and prenatal sex ratio. Since group size and sex ratio were highly correlated within a year (group size: Pearson’s r = 0.95; sex ratio: Pearson’s r = 0.80), we only included prenatal group size and sex ratio in the final model. All two-way interactions were included in the model, but later removed if their effect was not significant ^89^. Breeding season of birth, season (long vs. short rains), social group, mother ID, and father ID were included as random effects. All random effects other than breeding season of birth had variance components equal to zero and were thus removed from the final model to facilitate the computation of 95% confidence intervals. This had no effect on the estimates of fixed effects^90^. Furthermore, to investigate the relationship between alloparental care provided by males prior to reaching one year of age and their subsequent reproductive tactic, we used a Chi-square test to compare alloparental status (ever alloparent/never alloparent) and reproductive tactic (dispersed/remained) for all natal males (N = 185).

Finally, we sought to understand the fitness consequences of the alternative reproductive tactics using data from males with known complete life histories. Although it was not possible to compare the fitness consequences of the alternative reproductive tactics adopted by males born from same social group (since dispersers rarely remain within the study population; only 3 males in 16 years), we compared access to reproductive opportunities, reproductive success, lifetime inclusive fitness, and survivorship of resident and immigrant males. Previous studies of cooperatively breeding birds have used the same approach ^58^. Access to reproductive opportunities was quantified as: (i) the likelihood of a male ever breeding in its lifetime (Chi-squared test) (N_Imm_ = 59, N_Res_ = 103); (ii) lifetime breeding effort, or the number of breeding attempts as a proportion of an individual’s adult lifespan (Mann-Whitney test with continuity correction) (N_Imm_ = 59, N_Res_ = 103); and (iii) for males that bred at least once (N_Imm_ = 30, N_Res_ = 29), the age at first breeding (Mann-Whitney test with continuity correction). To evaluate the likelihood of a male obtaining any reproductive success in their lifetimes, we modeled the effect of dispersal status on nest success (1 = succeeded, 0 = failed) using a GLMM with a binomial error structure and “ logit” link function (N_succeeded_ = 49, N_Failed_ = 229). Since some males nested multiple times in their lifetimes, we included individual ID as a random effect in the model along with social group and breeding season.

Following Green and Hatchwell ^58^, we calculated lifetime inclusive fitness (I) as the sum of lifetime direct and indirect fitness according to the equation

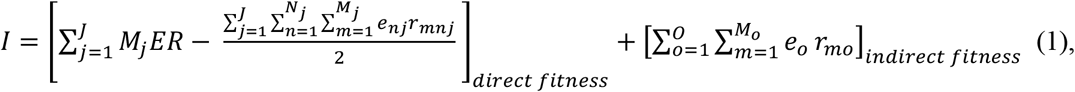

where direct fitness was calculated as the product of the number of fledglings per nest (M_j_), paternal care effort (E = 0.5, the other half being attributed to the mother), and the mean offspring-father relatedness (R = 0.5) summed over all successful nests (J), minus half of the indirect fitness attributed to alloparents (the other half being subtracted from the mother’s direct fitness). Indirect fitness attributed to alloparents at each nest was calculated as the product of the mean alloparental effort at a nest per social group per breeding season (e_nj_) and the alloparent’s relatedness to the fledglings (r_mnj_), summed over all fledglings (M_j_) and alloparents at the nest (N_j_) for all successful nests (J). Alloparents with r_mnj_ ≤ 0 received no indirect fitness. Indirect fitness of the focal male was similarly calculated as the product of the mean alloparental effort (e_o_) and its relatedness to the fledglings (r_mo_), summed over all of the nests he visited in his lifetime (O). We used population means instead of individual measures of alloparental effort because total observation time for nests varied. Alloparental effort was calculated as the proportion of time an individual spent attending a nest (both guarding and bringing food) relative to the length of the observation period ^41^.

We compared the lifetime inclusive, direct, and indirect fitness of resident and immigrant males using a Mann-Whitney test with continuity correction (N_Imm_ = 53, N_Res_ = 99). Since males accrue indirect fitness as juveniles, we excluded males born in 2001 short rains, the earliest breeding season in our dataset, during which limited focal observations were conducted at nests. To account for indirect fitness accrued by immigrant males in their natal group prior to dispersing, we additionally (i) compared fitness of residents and immigrants after excluding fitness accrued by residents during the first year of their life (see *SI*), and (ii) compared the fitness accrued in the first year of life by natal males that remained in their natal group to that accrued by natal males that subsequently dispersed (see *SI*). To account for mortality during dispersal, we conducted a sensitivity analysis by adding incremental proportions of immigrants with zero lifetime fitness to the dataset (see *SI*).

To determine whether differences in reproductive access and success between resident and immigrant males affected their lifetime inclusive fitness, we used GLMMs to model the effect of lifetime breeding effort and dispersal status on lifetime inclusive fitness of males. The first model had a binomial response (zero/nonzero lifetime inclusive fitness) and a “ logit” link function (N_Imm_ = 53, N_Res_ = 99). The second model had a continuous dependent variable of all nonzero inclusive fitness observations (N_Imm_ = 20, N_res_ = 47) with an “ identity” link function and a gamma error distribution. The fixed effects for both models were lifetime breeding effort and dispersal status. Social group was included as a random effect, but later removed from the second model, where its variance component equaled zero, to facilitate the computation of 95% confidence intervals ^90^.

We also built a GLMM to investigate the lifetime fitness consequences of adopting the tactic not predicted by the prevailing environmental conditions, with a binomial response variable (zero/nonzero lifetime inclusive fitness) and prenatal rainfall and dispersal status as fixed effects (N_Imm_ = 53, N_Res_ = 99). Although we did not have precise rainfall data from the immigrants’ natal sites, we used monthly rainfall data from three sites within the dispersal radius (∼30 km) (Shah and Rubenstein, unpubl. data) of our study population (Mpala Research Centre, MRC)—UHURU Central (12 km from MRC) ^91^, UHURU North (20 km from MRC) ^91^, and Nanyuki (38 km from MRC) (East African Livestock Early Warning System)—to demonstrate high spatiotemporal correlation of prenatal rainfall for immigrant and resident males (Table S1). Nonetheless, to take a more conservative approach, we used a categorical measure based on mean long-term pre-breeding rainfall at Mpala Research Centre (low prenatal rainfall: rainfall ≤ long-term mean; high prenatal rainfall: rainfall > long-term mean). Rainfall within 10 mm of the long-term mean was categorized as “ low” to account for the accuracy of the rainfall gauge ^87^.

Finally, we used a time-varying Cox proportional hazard model to determine whether dispersal status affected male survival (N_Imm_ = 59, N_Res_ = 103) ^42^. The model was built using the R package *survival* ^92^. We checked that our dataset did not violate the proportional hazard assumption using the “ cox.zph” function. Social group was included as a random factor. We performed all data analysis in R ^93^. VIF < 2 for all fixed effects in GLMMs, excluding interaction terms ^94^.

## Data Availability

Data and R code will be made available on Dryad.

## Acknowledgements

We thank B. Heidinger, M. Uriarte, M. VanAcker, J. Jensen, P. Downing, S. Caro, S. Siller, Y. Cheng, A. Earl, J. Guan, S. Guindre-Parker, Y. Haba, E. Greig, R. Gloag, M. Paquet, and an anonymous reviewer for statistical advice and comments on the manuscript. We are grateful to W. Watetu, G. Manyaas and J. Mosiany for their assistance in the field, as well as all other volunteers, students and field assistants that have contributed to this project since its inception in 2001.

