## Supplementary Information for "Prenatal environmental conditions underlie alternative reproductive tactics that drive the formation of a mixed-kin cooperative society"

This PDF file includes:

Supplementary Information Text
Figures S1 to S4
Table S1 to S2
SI References

**Supplementary Information Text**

**Estimating the age of dispersal for males**

We inferred the age of dispersal of males indirectly using four lines of evidence: (i) the modal age at which males disappear from their natal group (N = 155); (ii) the categorical age of immigrant males dispersing into the study population as determined by plumage and eye color (fledglings: ≤4 mo, sub-adults: 4 mo – 1 yr, adults: ≥1 yr) (N = 97); (iii) the likelihood of males being detected in the census in their first year of life (N = 103); and (iv) the minimum age of first breeding by resident males (N = 29). We found that (i) 33% of natal males that disappeared did so within the first six months of hatching (Fig. S2A). If six months is the modal age of dispersal, most immigrants into the study population should be in juvenile (<1 year of age; fledgling or sub-adult) plumage. However, (ii) 90% of male immigrants into the study population were in adult plumage (i.e., ≥1 year of age) (Fig. S2B). Depending on the month and season (short vs. long rains) of their birth, males would be between 8-11 months old at the start of the breeding season in which they reach sexual maturity. Coupled with individual variation in plumage and eye color, this may explain the distribution of categorical ages of immigrants into our study population. Thus, we reasoned that natal males disperse around 1 year of age but the majority of them are likely missed in the census during the one breeding season they spend in their natal group before they disperse, largely because of our methods for performing censuses. However, dispersal patterns remain to be rigorously quantified in superb starlings, which may be possible in the future with the advent of novel, lightweight GPS tracking devices.

Since our census effort is mostly opportunistic, it is common for individuals to be missed for multiple breeding seasons before being seen again, especially if they did not breed or act as an alloparent [42]. We found that (iii) of all the males that remained in their natal group past 1 year of age, 64% were missed in the census at age 0.5 years (six months). Furthermore, the likelihood of missing individuals was higher in the period preceding the short rains breeding seasons (70%) than in the period preceding the long rains breeding seasons (48%), which directly matches the variation in census effort. Finally, when we set age of dispersal as 1 year, 52 of 185 natal males (28%) were classified as "dispersed" in our dataset, a proportion that closely matches the proportion of male immigrants in the study population (see *Results*) and gives us more confidence in our inference of dispersal age. Moreover, (iv) the minimum age of first breeding by resident males was at one and a half years old, further suggesting that males are reproductively immature in their first year of life and may thus disperse at around one year of age. While we cannot confirm that males dispersed, not died, the increased likelihood of dispersal following benign prenatal conditions (see *Results*) suggests that these males did indeed disperse, since mortality would be higher following harsher conditions. To further to ensure that this relationship is not driven by mortality, we built a linear model of hatchling mass pre-fledging (at ten days-old, modal age at banding) and prenatal rainfall. Social group, breeding season, and nest ID were included as random effects. Hatchling mass did not vary significantly with variation in prenatal rainfall ($N = 47$, $Z = 1.19$, $P = 0.25$), unlike other studies where the effect of harsh ecological conditions on post-fledging survival has been shown to be mediated by hatchling mass pre-fledging (Bourne et al., 2020; Ridley et al., 2021).

**Calculating additional measures of lifetime fitness**

When the fitness accrued by natal males in their first year was excluded, residents and immigrant males had similar indirect and inclusive lifetime fitness (indirect: $U = 2607$, $P = 0.94$, inclusive: $U = 2765$, $P = 0.52$). Since males do not accrue any direct fitness as juveniles, excluding fitness gained during the first year did not affect variation in direct fitness by dispersal status. Natal males that dispersed accrued significantly less fitness in their first year than natal males that remained in their natal group, suggesting that immigrants into our study population did not accrue considerable fitness prior to dispersing ($N = 185$, $U = 4241$, $P = 0.003$). Furthermore, acting as an alloparent during the juvenile stage may be part of the tactic of philopatry as has been shown in other cooperatively breeding species (Clutton-Brock et al. 2002, Green and Hatchwell 2018).

To account for mortality during dispersal, we added immigrants with zero lifetime fitness to the dataset in 10% increments and compared the lifetime fitness of immigrants and residents. When comparing only fitness accrued as an adult (i.e., excluding fitness accrued by residents in their first year of life), we found that residents had significantly higher direct, indirect, and inclusive lifetime fitness when 70%, 40%, and 50% of immigrants with zero fitness were added, respectively (Table S1). With fitness from the first year of the residents' life included, the inclusive fitness of residents was significantly higher with 20% of immigrants with zero fitness added (Table S1). While we do not have an estimate of mortality during dispersal in our study population, we have no evidence of floaters in this species, suggesting that mortality rate of dispersers is much lower than estimates in other cooperatively breeding species (*e.g.*, Kingma et al. 2016, Walters et al. 1992).

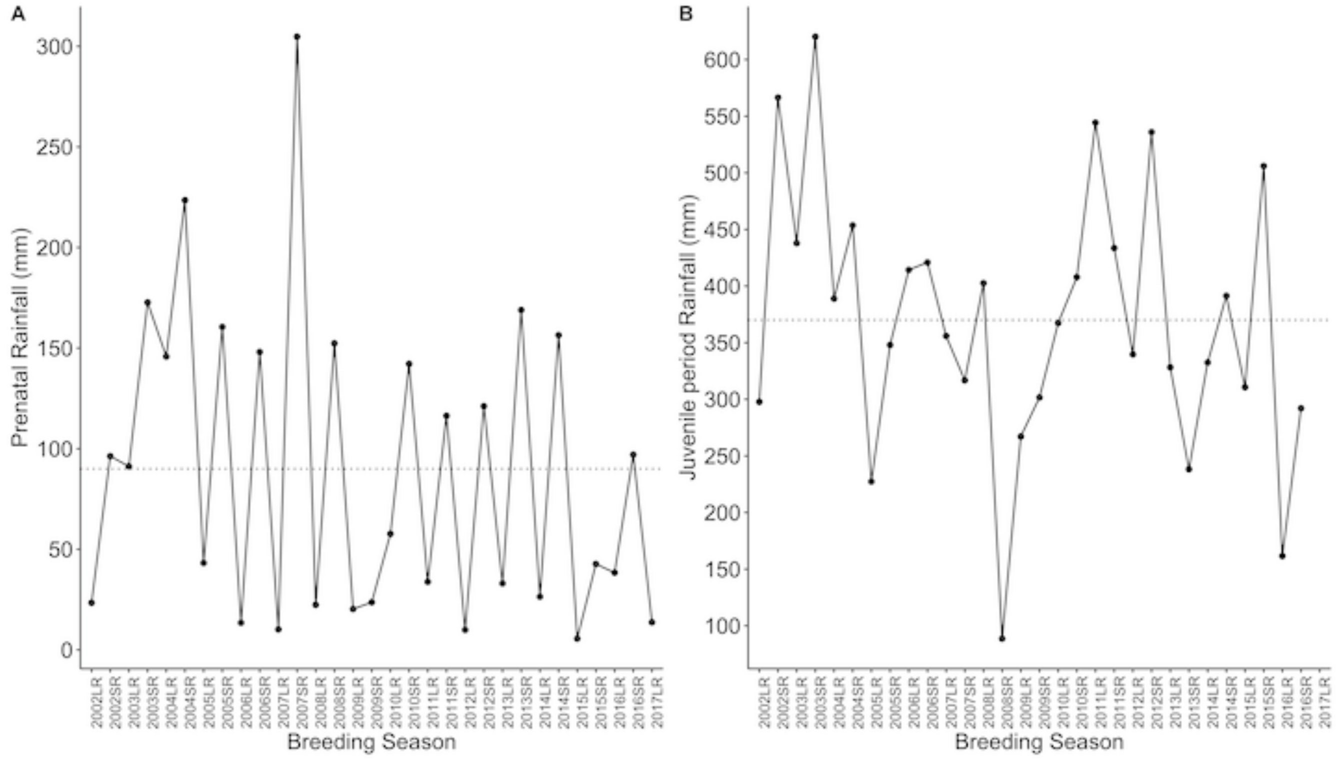


**Figure S1. Variation in rainfall during the study period.** Superb starlings in the study population breed twice a year in the long (LR) and short (SR) breeding seasons. (A) Variation in prenatal rainfall during the study period. (B) Variation in juvenile stage rainfall during the study period based on the breeding season in which an individual was born. Dotted lines indicate the mean.

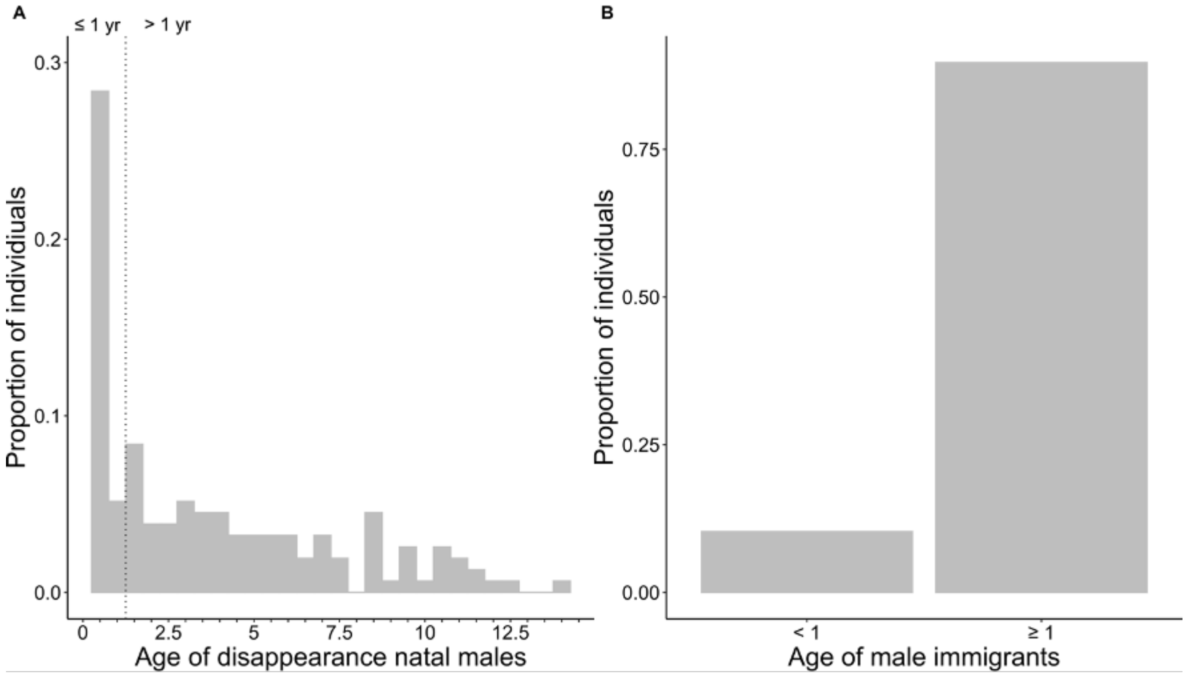


**Figure S2. Evidence for age of dispersal of male superb starlings.** (A) The modal age of disappearance of natal males is six months (N = 155). Age is in 0.5-year (i.e., six month) increments. (B) However, the majority of immigrant males are caught in adult plumage (≥1 year of age) (N = 97). Thus, we concluded that males disperse around 1 year of age and are likely missed in the census during the one breeding season they spend in their natal group before dispersing.

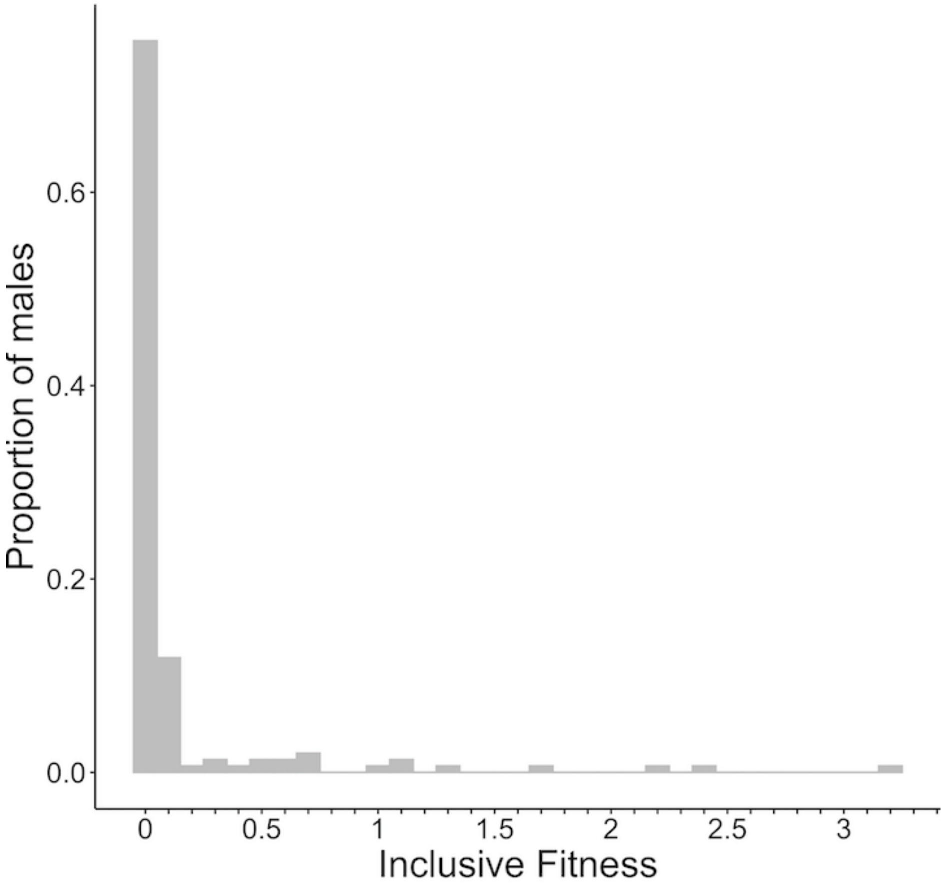


**Figure S3. Lifetime inclusive fitness of male superb starlings.** Histogram show the distribution of lifetime inclusive fitness of males (N = 152).

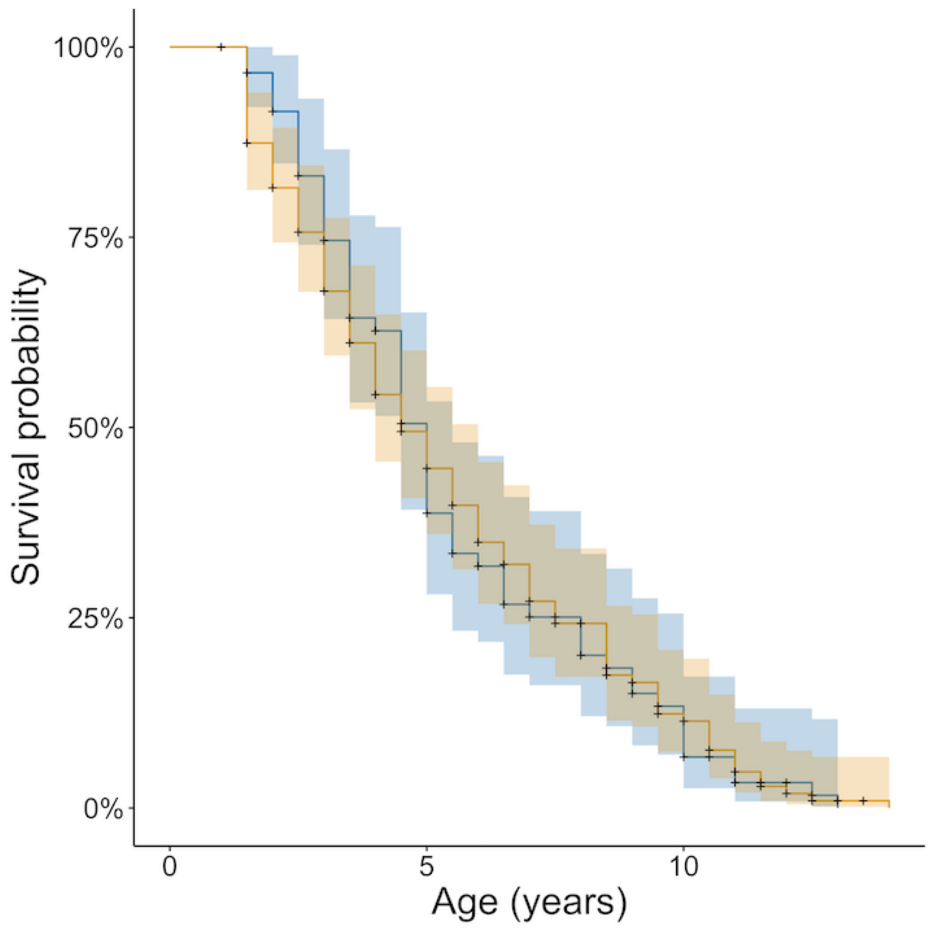


**Figure S4. Survivorship of resident and immigrant male superb starlings.** Male survival did not differ by dispersal status (immigrants = blue, residents = orange) ($N_{Imm}$ = 59, $N_{Res}$ = 103, P = 0.95). Model estimates (solid lines) are bound by 95% confidence intervals.

**Table S1. Sensitivity analysis accounting for potential mortality of immigrant male superb starlings during dispersal.** Results of adding immigrants with zero fitness in increments of 10% to analysis of variation in fitness by dispersal status. Fitness of immigrant and resident males was compared using Mann-Whitney tests. We compared all categories of lifetime fitness found to be similar for residents and immigrants (i.e., direct fitness, inclusive fitness, indirect fitness excluding fitness gained in first year of life by residents, and inclusive fitness excluding fitness gained in first year of life by residents) (see *Results*). Bold indicates threshold at which fitness for immigrants and residents becomes significantly different.

| Fitness | Percentage of immigrants with zero fitness added | U | p-value |
|---|---|---|---|
| Direct fitness | 10 | 2968 | 0.75 |
| | 20 | 3279 | 0.94 |
| | 30 | 3724 | 0.83 |
| | 40 | 4258 | 0.60 |
| | 50 | 5059 | 0.36 |
| | 60 | 6216 | 0.16 |
| | **70** | **8219** | **0.04** |
| Inclusive fitness | 10 | 2493 | 0.09 |
| | **20** | **2675** | **0.03** |
| Indirect fitness *excluding first year for residents* | 10 | 2811 | 0.63 |
| | 20 | 3049 | 0.37 |
| | 30 | 3389 | 0.15 |
| | 40 | 3797 | 0.05 |
| | **50** | **4409** | **0.01** |
| Inclusive fitness *excluding first year for residents* | 10 | 2696 | 0.83 |
| | 20 | 3207 | 0.81 |
| | 30 | 3547 | 0.42 |
| | 40 | 3955 | 0.17 |
| | **50** | **4567** | **0.03** |

**Table S2. Correlation in rainfall between sites within the dispersal radius of the study site.** Numbers represent Pearson correlation coefficients (r) of monthly rainfall (2008-2017; N = 103) between three sites within the approximate dispersal radius of the study site, Mpala Research Centre (MRC), from which we could obtain rainfall data: UHURU Central (12 km), UHURU North (20 km), and Nanyuki (38 km).

| | MRC | UHURU Central | UHURU North | Nanyuki |
|---|---|---|---|---|
| MRC | - | - | - | - |
| UHURU Central | 0.85 | - | - | - |
| UHURU North | 0.82 | 0.87 | - | - |
| Nanyuki | 0.78 | 0.73 | 0.70 | - |
